# Effect of extracorporeal shock wave therapy on equine umbilical cord blood mesenchymal stromal cells *in vitro*

**DOI:** 10.1101/2020.01.10.901439

**Authors:** Ramés Salcedo-Jiménez, Judith Koenig, Olivia Lee, Thomas W.G. Gibson, Pavneesh Madan, Thomas G. Koch

## Abstract

Extracorporeal shock wave therapy (ESWT) has been shown to induce different biological effects on a variety of cells, including regulation and stimulation of their function and metabolism. ESWT can promote different biological responses such as proliferation, migration, and regenerations of cells. Recent studies have shown that mesenchymal stromal cells (MSCs) secrete factors that enhance the regeneration of tissues, stimulate proliferation and differentiation of cells and decrease inflammatory and immune-reactions. Clinically, the combination of these two therapies has been used as a treatment for tendon and ligament lesions in horses; however, there is no scientific evidence supporting this combination of therapies *in vivo*. Therefore, the objectives of the study were to evaluate the effects of ESWT on equine umbilical cord blood mesenchymal stromal cells (CB-MSCs) proliferative, metabolic, migrative, differentiation, and immunomodulatory properties *in vitro.* Three equine CB-MSC cultures from independent donors were treated using an electrohydraulic shock wave generator attached to a water bath. All experiments were performed as triplicates. Proliferation, viability, migration and immunomodulatory properties of the cells were evaluated. Equine CB-MSCs were induced to evaluate their trilineage differentiation potential. ESWT treated cells had increased metabolic activity, showed positive adipogenic, osteogenic, and chondrogenic differentiation, and showed higher potential for differentiation towards the adipogenic and osteogenic cell fates. ESWT treated cells showed similar immunomodulatory properties to none-ESWT treated cells. Equine CB-MSCs are responsive to ESWT treatment and showed increased metabolic, adipogenic and osteogenic activity, but unaltered immunosuppressive properties. *In vivo* studies are warranted to determine if synergistic effects occur in the treatment of musculoskeletal injuries if ESWT and equine CB-MSC therapies are combined.

## Introduction

Extracorporeal shock wave therapy (ESWT) is one of the leading treatments of certain orthopedic diseases in humans such as plantar fasciitis and lateral epicondylitis and more recently it has been used to treat Achilles and patellar tendinopathies, [1,2]. ESWT is also a common treatment for tendon and ligament injuries in the horse particularly suspensory ligament desmitis [3]. At present, it is also used in horses for osteoarthritis when other treatments are ineffective [4]. ESWT uses acoustic waves generated mechanically outside of the body that can be focused to a specific point within the body. Shock waves used for medical purpose are generated in a fluid medium, by one of three different generators: electrohydraulic, piezoelectric or electromagnetic. The shape of the acoustic wave is characterized by an initial positive rapid phase, of high amplitude, followed, by a sudden phase of mild negative pressure, and then returns to the ambient values. Their peak pressure is high - up to 100 mpa (500 bar) with a rapid rise (<10 ns) in pressure, of short duration (<10 ms) and a broad range of frequency [5,6].

A variety of cells in culture, including MSCs, have shown to be responsive to ESWT. Human bone marrow stromal cells (hBMSCs) showed increased proliferation, migration and the rate of apoptosis activation was reduced after treatment with focused ESWT, and all treated cells maintained their differentiation potentials [7]. Rat adipose-derived stromal cells (ASCs) responded to ESWT with an elevation of mesenchymal markers and a higher capacity to differentiate towards adipogenic and osteogenic lineages with minimal changes in proliferation [8]. When equine adipose tissue-derived mesenchymal stromal cells were treated with ESWT, they showed increased proliferation, but no effects were observed in their differentiation potential [9]. The increased proliferation after shockwave treatment may be mediated through purinergic receptors via downstream ErK1/2 signals following activation by extracellular ATP [10].

MSCs are multipotent cells capable of differentiation into osteogenic, adipogenic and chondrogenic lineages [11]. Bone marrow aspirates and adipose tissue are the most studied and used sources to obtain MSCs for clinical cases in the horse [12]. It has been suggested that MSCs from umbilical cord blood (CB-MSCs) might have superior immune tolerance, proliferative potential, and differentiation potency [13]. MSCs may function in two different ways, first in a progenitor function by direct tissue integration and second in a non-progenitor function through secretory products that have trophic and immunosuppressive effects [14]. Recent studies have shown that MSCs secrete factors that enhance regeneration of injured tissue, stimulate proliferation and differentiation of endogenous cells and decrease inflammatory and immune reactions [14–16].

Clinically, the combination of ESWT and MSC therapies have been used as a treatment for tendon and ligament lesions in horses without scientific evidence supporting this approach. We hypothesized that ESWT will affect the progenitor functions and non-progenitor functions of equine CB-MSCs. The aim of this study was to evaluate if ESWT affects equine CB-MSCs proliferative, metabolic, migrative, differentiation, and immunomodulatory properties *in vitro*.

## Material and methods

### Culture of equine CB–MSCs and *in vitro* extracorporeal shock wave treatment

The equine CB-MSC were established as previously described by Koch *et al.* [17]. Briefly, the mononuclear cell fraction (MNCF) was cultured and non-adherent cells were removed through successive complete medium changes. When cell numbers allowed, the cells were passaged and expanded until they were cryopreserved for later use.

Three cryovials of equine CB-MSC isolated from different donors were used in these studies. The cryovials had been stored in liquid nitrogen from 1 to 9 months prior to the study and all contained cells at passage 3. The cells were thawed, seeded at a cell density of 5,000 cells/cm^2^ culture surface area and expanded until ∼80% confluence to obtain sufficient cell numbers for all the experiments and assays of these studies.

The *in vitro* shock wave treatment was performed as described by Holfeld *et al*. using an electrohydraulic shock wave generator (VersaTron, Pulse Veterinary Technologies, Alpharetta GA, USA) [18]. Equine CB-MSCs were treated at a confluence of 90% in T-25 cell culture flasks. The flasks were placed in front of the applicator at a distance of 5 cm from the probe to the cell layer inside the water container in direct contact with pre-warmed (37°C) water. The focused shock waves were applied with the following parameters: an energy flux density of 0.1 mJ/mm^2^, frequency of 3Hz and 300 impulses (Fig 1).

**Fig 1.**
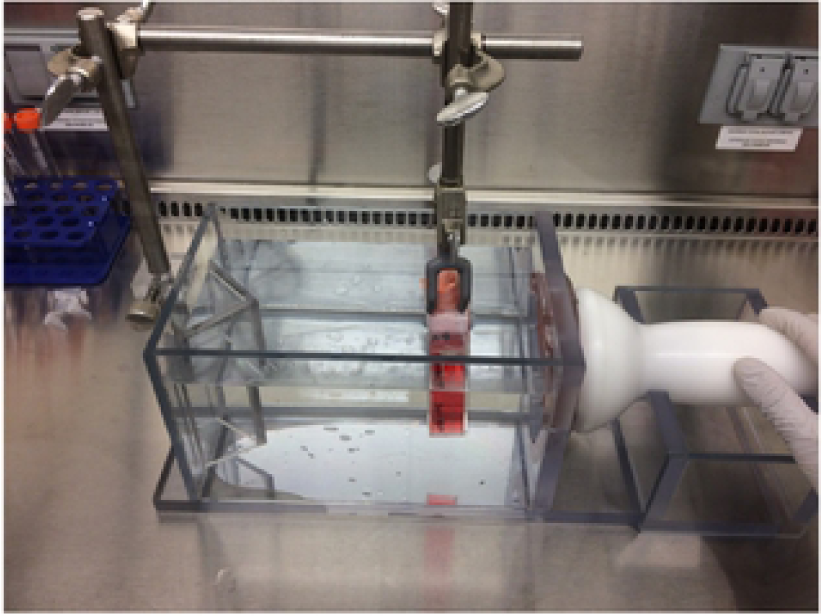
Water bath for ESWT application. The bottom of the culture flask containing the adherent cells are facing the ESWT probe.

The control group was maintained in the same culture conditions, consisting of equilibrated expansion media or differentiation media without shock wave exposure. All cell culture experiments were performed as triplicates from 3 different donors. The process is shown in flow the chart (Fig 2).

**Fig 2.**
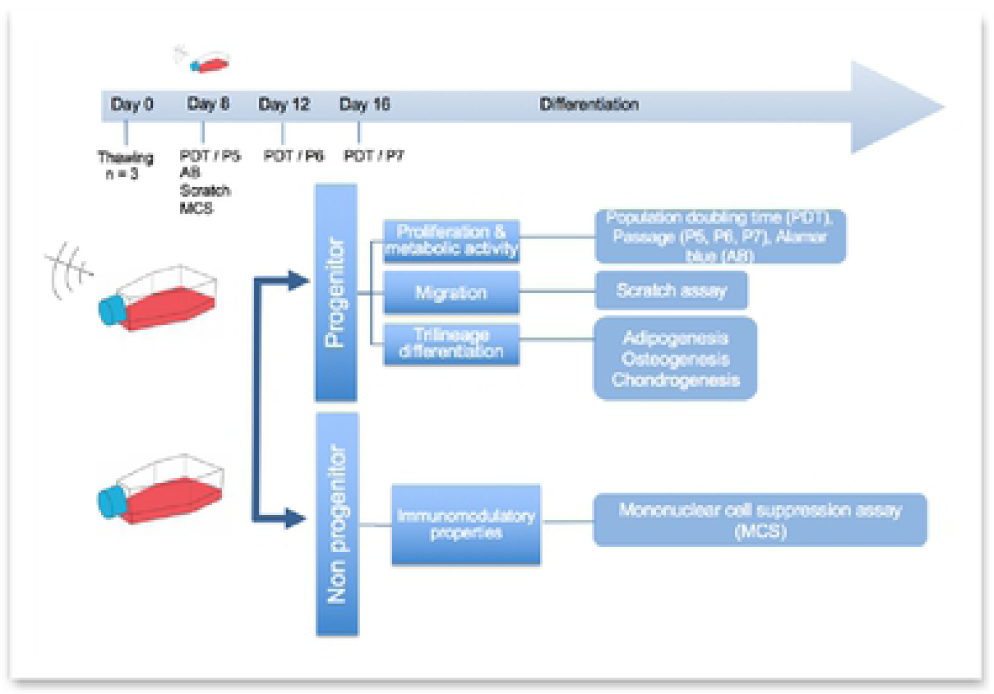
Experimental design. Time points for ESWT, respective passage numbers (P5, P6, P7) and population doubling time (PDT). Alamar blue assay (AB). scratch (migration) assay and mononuclear cell suppression assay (MCS). Differentiation was initiated after treatment on day 8.

### Proliferation

Cells were detached using trypsin 0.025% (Lonza, Walkersville, MD, USA) and re-seeded in 6 well culture flasks. To document cell morphology, digital images were obtained prior to detachment at passage 5, 6 and 7 using phase-contrast microscopy and Q-Capture software (Q-Imaging, Surrey, BC, Canada).

Doubling times for passage 5, 6 and 7 were calculated according to Tessier *et al*. :

1.Cell–doubling number (CD) = In(Nh/Ns)/In 2

In = Natural Logarithm

Nh = harvest cell number

Ns = seed cell number

2. Cell doubling time (DT) = CT/CD

CT = cell culture time [19].

### Metabolic activity

Equine CB-MSC were seeded in 96-well plates at 5,000 cells/cm^2^ in 300 μl of culture media. The following day the culture media was changed. On day 4, the culture media was replaced with 300 μl of 10% resazurin (Sigma Aldrich, Oakville, ON, Canada) in phosphate-buffered saline (PBS), incubated at 37°C for 4 h protected from light and read by a plate reader (Spectramax i3, Molecular Devices, Sunnyvale, CA, USA) at 585 nm using an excitation wavelength of 555 nm.

### Migration (scratch assay)

For the scratch migration assay, equine CB-MSC were seeded at 4 × 10^4^ cells in 70 μl into each dish (Culture-Insert 2 well in μ-Dish 35mm, high Ibidi cat# 80206) and incubated, cells were grown to confluence. The Culture-Insert 2 well was removed, and the cell layer was washed with PBS to remove debris and 2 ml of culture medium were added. Images were taken at 0, 12, 24, 36, and 48-hour time points or until complete closure of the scratch wound was observed using a motorized inverted microscope (IX81, Olympus America Inc., Melville, NY, USA). The dishes were incubated between each time point. After all images were acquired, measurements of the surface area of each scratch at each time points were performed with Photoshop (Adobe Systems Incorporated, NY, USA).

### Trilineage differentiation studies

For adipogenesis, equine CB-MSC were seeded at 2.1 × 10^4^ cells/ml in 6 well plates, media was changed every 2-3 days. After the cells reached 100% confluence the cells were exposed to adipogenic induction media (BulletKit, Adipogenesis, Lonza). The induction medium consisted of 1 mM dexamethasone, 0.5 mM 3-isobutyl-1-methyl-xanthine (IBMX), 10 mg/ml recombinant human (rh) insulin, 0.2 mM indomethacin and 10% fetal calf serum (FCS) in DMEM. Media was changed every 2-3 days as previously described, Koch *et al.* [17]. The adipogenic potential was assessed after 14 days of exposure to induction media by Oil Red O staining to observe lipid droplets and AdipoRed™ assay (Lonza).

For Oil Red O staining, all the media was aspirated off the wells, wash once with 2 ml of PBS, then replaced with 2 ml of 10% formalin (Fischer Scientific, Whitby, ON, Canada) for 30 min at room temperature to fixate the cells, the formalin was replaced with 2 ml of sterile water (Milli-Q water, Millipore, Mississauga, ON, Canada) for 2 minutes and then replaced with 60% isopropanol (Fischer Scientific) for 5 minutes, the isopropanol was replaced with 2 ml of Oil Red O (Sigma-Aldrich,) working solution [3 parts (300 mg Oil Red O in 100 ml of 99%o isopropanol) mixed with 2 parts sterile water], after 5 minutes the Oil red O solution was washed of indirectly with tap water. Two ml of Harris hematoxylin (Sigma Aldrich) were added to the wells for 1 minute before being aspirated and the wells washed with warm tap water.

The AdipoRed Assay (Lonza) was performed according to the manufacturer’s protocol for 96 well plates. Briefly, equine CB-MSC were induced for 14 days. The supernatant was removed and rinsed with 200 μl PBS. Then each well was filled with 200 μl of PBS, 5 ul of reagent were injected followed by shaking for 1 second. The fluorescence was measured with excitation at 485 nm and emission at 520 nm as previously described by Koch *et al*. [17].

Chondrogenesis differentiation was performed in pellet culture as previously described by Co *et al*. [20]. Approximately 2.5 × 10^5^ CB-MSCs were suspended in 200 μl of the chondrogenic medium, containing High Glucose Dulbecco’s modified Eagle medium (HG-DMEM) (Sigma Aldrich; Oakville, ON), 200 mM Glutamax (Invitrogen; Burlington, ON, Canada), 100 mM sodium pyruvate (Invitrogen), 1 ABAM (Invitrogen), 0.1 mM dexamethasone (Sigma Aldrich), 100 mg/ml ascorbic acid-2 phosphate (Sigma Aldrich), 40 mg/ml proline (Sigma Aldrich), 1 ITS (BD Biosciences; Mississauga, ON, Canada), and 10 ng/ml TGF-β3 (R&D Systems; Minneapolis, MN, USA), the cells were plated in V-bottom polypropylene 96 well plates (Phenix #MPG-651201) and centrifuged at 200x g for 10 min and incubated for 3 weeks. The media was changed every 2-3 days. Chondrogenic potential was assessed by Toluidine Blue to visualize glycosaminoglycan (GAG) and biochemistry.

For Toluidine blue stain, the pellets were fixed in 10% formalin, dehydrated in isopropanol and embedded in paraffin and sectioned (5 uM). Slides were deparaffinized and rehydrated prior to staining. The stock solution of Toluidine blue contained 1 g powder (Sigma Aldrich) in 100 ml 70% EtOH diluted to 10 % with 1% NaCl. Sections were stained in Toluidine blue working solution for 2 min and washed in three changes of distilled water.

For biochemistry evaluation, the pellets were digested in 40 mg/ml papain (Invitrogen; Burlington, ON, Canada), 20 mM ammonium acetate, 1 mM EDTA, and 1 mM dithiothreitol (DTT) (Sigma Aldrich) for 48 h at 65°C vortexing at 24 h. Stored at – 40°C until further analysis. The proteoglycan content of the digest was estimated by quantifying the amount of sulfated glycosaminoglycans (GAG), using the dimethylmethylene blue dye and spectrophotometry with a wavelength of 525 nm. The standard curve for the proteoglycan content assay was generated using chondroitin sulfate (Sigma Aldrich). DNA content was assessed using a commercial kit: DNA quantitation kit e Fluorescence assay (Sigma Aldrich) in a 96-well plate. The assay was carried out according to the manufacturer’s instructions.

For osteogenesis, equine CB-MSC were seeded at a density of 3,000 cells/cm^2^ in 6 well plates, media was changed every 2-3 days. After the cells reached 100% confluency the cells were exposed to osteogenic induction media (BulletKit, osteogenic, Lonza). The induction medium consisted of DMEM-LG, 10% FBS, 1% L-glutamine, 1% antibiotic-antimycotic (ABAM), 100 nM dexamethasone, 10 mM β-glycerophosphate, and 0.05 mM ascorbic acid-2-phosphate. The media was changed every 3-4 days. The osteogenic potential was assessed by Alizarin Red S staining for calcium deposition and semi-quantitative alkaline phosphatase enzyme activity assay.

For Alizarin Red S staining, the pH of the Alizarin solution was adjusted to 4.1-4.3 with ammonium hydroxide. The media was aspirated from each well and the cells were fixed in ice-cold 70% ethanol for 5 min at room temperature. Alcohol was aspirated and the wells were rinsed twice (5 min) with water, the water was aspirated, and 1 ml of 2% Alizarin Red S solution was added, it was incubated for 3 minutes at room temperature and removed, wells were washed 5 times with 2 ml of water.

For Alkaline phosphatase enzyme activity assay, the culture wells were rinsed twice with PBS, 0.5 ml of PBS was added to the well and the cells were scraped off and transferred to a 1.5 ml Eppendorf tube. The wells were rinsed with 0.5 ml PBS, which was transferred to the Eppendorf tube as well. The sample was centrifuged at 5000 Å∼ g for 8 seconds. All the PBS was aspirated, and the cell pellet was resuspended in 0.1 ml lysis buffer by pipetting the cells up and down several times. The lysis buffer consisted of (500 ml lysis buffer: 250 mg sodium deoxycholate, 5 mg phenylmethyl sulfonyl fluoride (PMSF), 5 mg aprotinin, 500 ml nonidet P-40, 500 ml 10% (wt/vol) SDS). The Eppendorf tubes were left on ice for 5 minutes and vortexed for 30 seconds, then they were microcentrifuged at maximum speed for 10 minutes at 6°C. Fifty microliters of the supernatant were added in triplicate from each osteo-induced and control sample to a well in a 96 well plate, then the kit, p-Nitrophenyl Phosphate Liquid Substrate System (Sigma-Aldrich), was used and 50 μl of p-nitrophenyl phosphate (pNPP) were added to each well. The absorbance was read at 405 nm on a microplate reader as soon as possible after adding the pNPP (time 0) and subsequent every 5 minutes for 20 minutes. The absorbance was read against 100 μl of undiluted pNPP. Covering it with aluminum foil screened the microplate from light between readings.

### Mononuclear cell suppression assay (MSA)

A pooled peripheral blood mononuclear cell (MNC) donor MSA was used to evaluate equine CB-MSCs for their ability to suppress MNC proliferation *in vitro.* Briefly, MNCs from five unrelated donors were mixed in equal ratios and stimulated with the mitogen concanavalin A (Sigma Aldrich). Negative control pooled MNCs were not stimulated with mitogen. Treated and non-treated CB-MSC cultures were irradiated (20 Gy) and mixed with stimulated pooled MNCs at a ratio of 1:10 (CB-MSC:MNC) in MNC media (RPMI 1640 with 10% horse serum, 1% L-glutamine and 1% pen/strep) and then seeded in 48 well plates. Reactions were incubated for four days at 38°C in 5% CO_2_. After five days of co-culturing, cells were stained with Bromodeoxyuridine (BrdU) and assessed with a BrdU ELISA kit (Roche, Mississauga, ON, Canada) following the manufacturer’s protocol.

## Statistical Analysis

For statistical analysis, the effect of ESWT delivered on equine CB-MSCs was evaluated. A statistical program (SAS 9.2, Proc MIXED, SAS Institute Inc., Cary, North Carolina, USA) was used to fit a general linear mixed model. For proliferation, the design was a two-factor factorial in a randomized complete block design with fixed effect factors treatment and passage. Interactions were also included in the model. For metabolic activity, the data was divided by a million for presentation. The design was also a two-factor factorial in a randomized complete block design with fixed effect factor cell culture and treatment. For migration, the design was a two-factor factorial in a randomized complete block design with fixed effect factors treatment and time. To accommodate time being a repeated measure, the following correlation structures (offered by SAS) were attempted: ar(1), arh(1), toep, toep(2)-toep(5), toeph toeph(2)-toeph(5) un un(2)-un(5), and one was chosen based on Akaike Information Criterion (AIC). For adipogenesis, the design was a two-factor factorial in a randomized complete block design by subsampling with fixed effect factors cell culture and treatment. And for osteogenesis to accommodate time being a repeated measure AIC was chosen. Repeated measures were nested in cell culture by treatment. Again, these error structures were attempted: ar(1) arh(1) toep toep(2)-toep(4) toeph toeph(2)-toeph(4) un un(2)-un(4). For the mononuclear cell suppression assay, a one-way ANOVA was performed with Tukey’s multiple comparisons test. Differences were considered significant at p ≤ 0.05.

To assess the ANOVA assumptions, comprehensive residual analyses were performed. The assumption of normality was formally tested by use of Shapiro-Wilk, Kolmogorov-Smirnov, Cramér-von Mises, and Anderson-Darling tests. Residuals were plotted against the predicted values and explanatory variables that were used in the models to look for outliers, bimodal distributions, the need for data transformations, or other issues that should be addressed.

## Results

### Proliferation

Treated and untreated equine CB-MSC showed an elongated, spindle-shaped fibroblast-like morphology. There were no obvious morphological differences between the treated and the untreated equine CB-MSCs.

One outlier was identified but was retained in the analysis. The data were transformed using log-based 2 and the assumption of the ANOVA analysis was adequately met and unequal variances in passages were accommodated. Population-doubling time was measured from passage 5 to 7. Mean (95% confidence interval (CI)) doubling time in days was as follows: passage 5, 1.67 days (0.83 – 3.14 days) for treated equine CB-MSC and 1.39 days (0.80 – 3.03 days) for untreated CB-MSC, for passage 6, 1.43 days (0.68 – 2.41 days) for treated equine CB-MSC and 1.46 days (0.67 – 2.39 days) for untreated equine CB-MSC and for passage 7, 2.42 days (1.11 – 4.30 days) for treated equine CB-MSC and 2.39 days (1.11 – 4.35 days) for untreated equine CB-MSC. There were no differences (*p > 0.05*) between treated and untreated equine CB-MSC (Fig 3).

**Fig 3.**
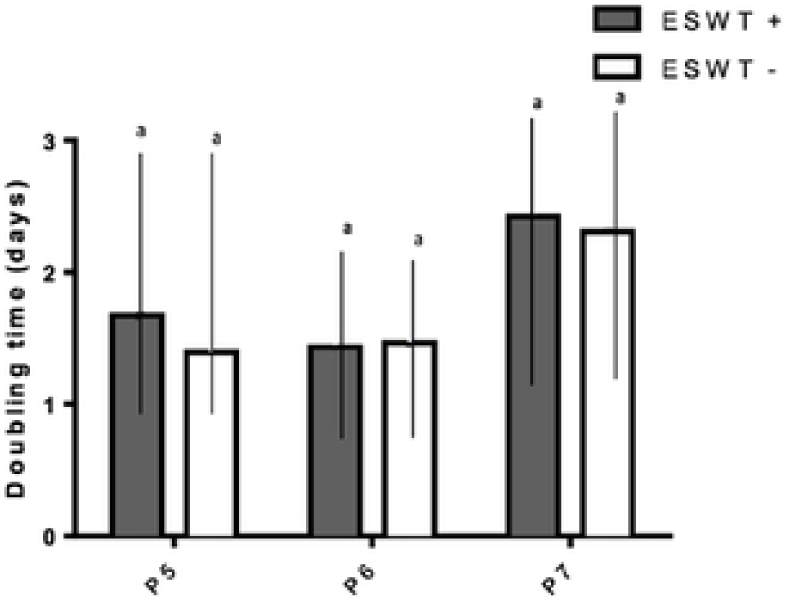
Proliferation. Population doubling times (days) for treated (ESWT +) and untreated (ESWT -) equine CB-MSCs for passage 5 (P5), 6 (P6) and 7 (P7). Results are shown as mean and CI 95% (*p* > *0.05*). No differences were observed.

### Metabolic activity

The data was divided by a million to allow for statistical analysis. Treated equine CB-MSC had higher metabolic activity than untreated equine CB-MSC as evaluated by the Alamar blue assay (p = 0.0002). Mean relative fluorescent units (RFU) was 102.29 (84.95 – 119.62 RFU) for treated equine CB-MSC and 65.10 RFU (47.76 – 82.43 RFU) for untreated equine CB-MSC (Fig 4).

**Fig 4.**
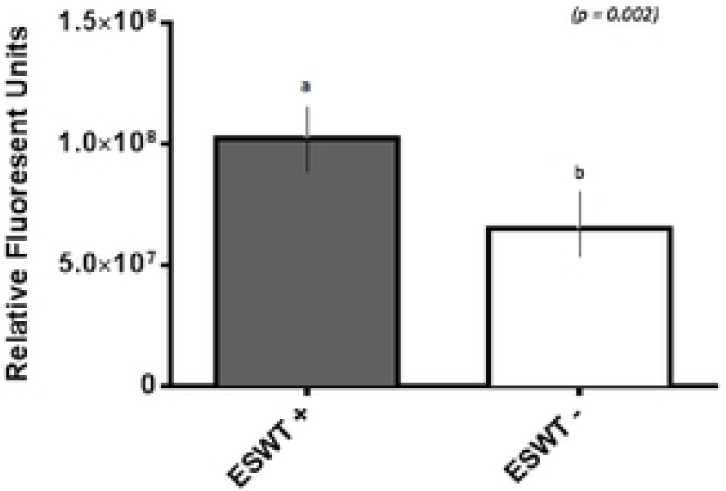
Metabolic activity. Relative fluorescent units for treated (ESWT +) and untreated (ESWT -) equine CB-MSCs. Results are shown as mean and CI 95% (*p* = *0.0002*) Values with different letters are statistically different.

### Migration (scratch assay)

Data was mildly non-normal, unequal variances in time were accommodated, and the assumptions were not well met. For the scratch assay, the results obtained at the 60-h time point were removed from the analysis as most of the scratches were already closed. For all the experiments, the time to closure among groups was similar. There was no difference for each area measured between the treated and the untreated equine CB-MSC (Fig 5).

**Fig 5.**
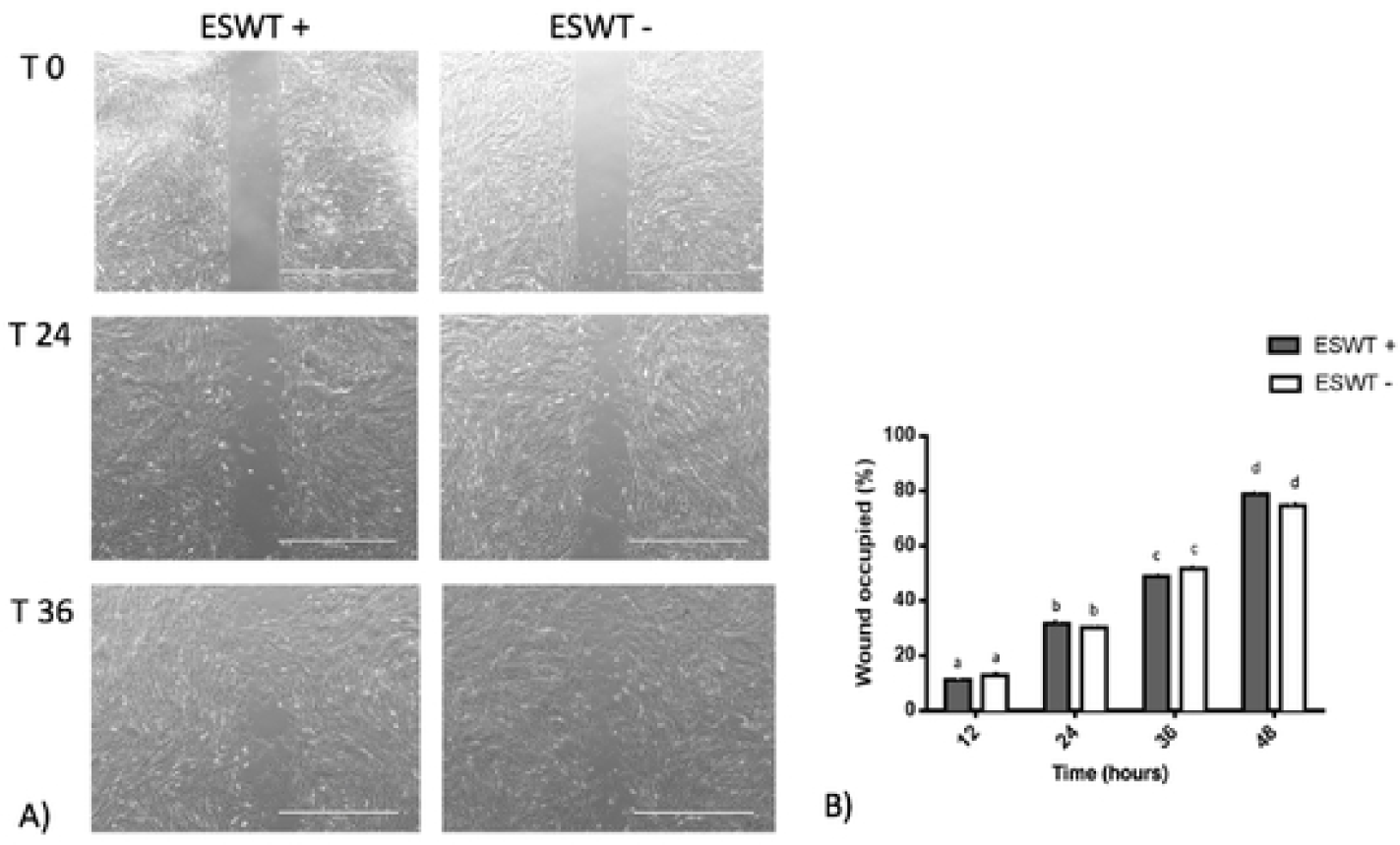
Migration. A) Representative scratch assay for treated (ESWT +) and untreated (ESWT -) equine CB-MSC. Images were obtained using a motorized inverted microscope until closure of the scratch in the cell layer. B) Percentage of wound occupied, calculated by dividing the none recovered area at 12, 24, 36 and 48 hours respectively by the initial wound area at 0 hours and subtracting this value as a percentage from 100%. There were no differences (*p* > *0.05)* between the treated (ESWT +) and untreated (ESWT -) equine CB-MSC at each time point.

### Trilineage differentiation

Positive trilineage potency was observed for the treated and the untreated equine CB-MSC, no subjective differences were observed for the differentiation capacities when evaluated by Oil Red O, Alizarin Red S, and Toluidine blue staining, respectively. Increased adipogenic differentiation occurred in the treated equine CB-MSC compared to the untreated cells based on a higher intracellular lipid formation (*p = 0.0002*). Higher osteogenic differentiation occurred in the treated equine CB-MSC compared to the untreated cells as demonstrated by a higher alkaline phosphatase activity in the initial minutes (p < 0.0001) (Fig 6). There were no differences for chondrogenesis as evaluated on biochemistry and wet mass.

**Fig 6.**
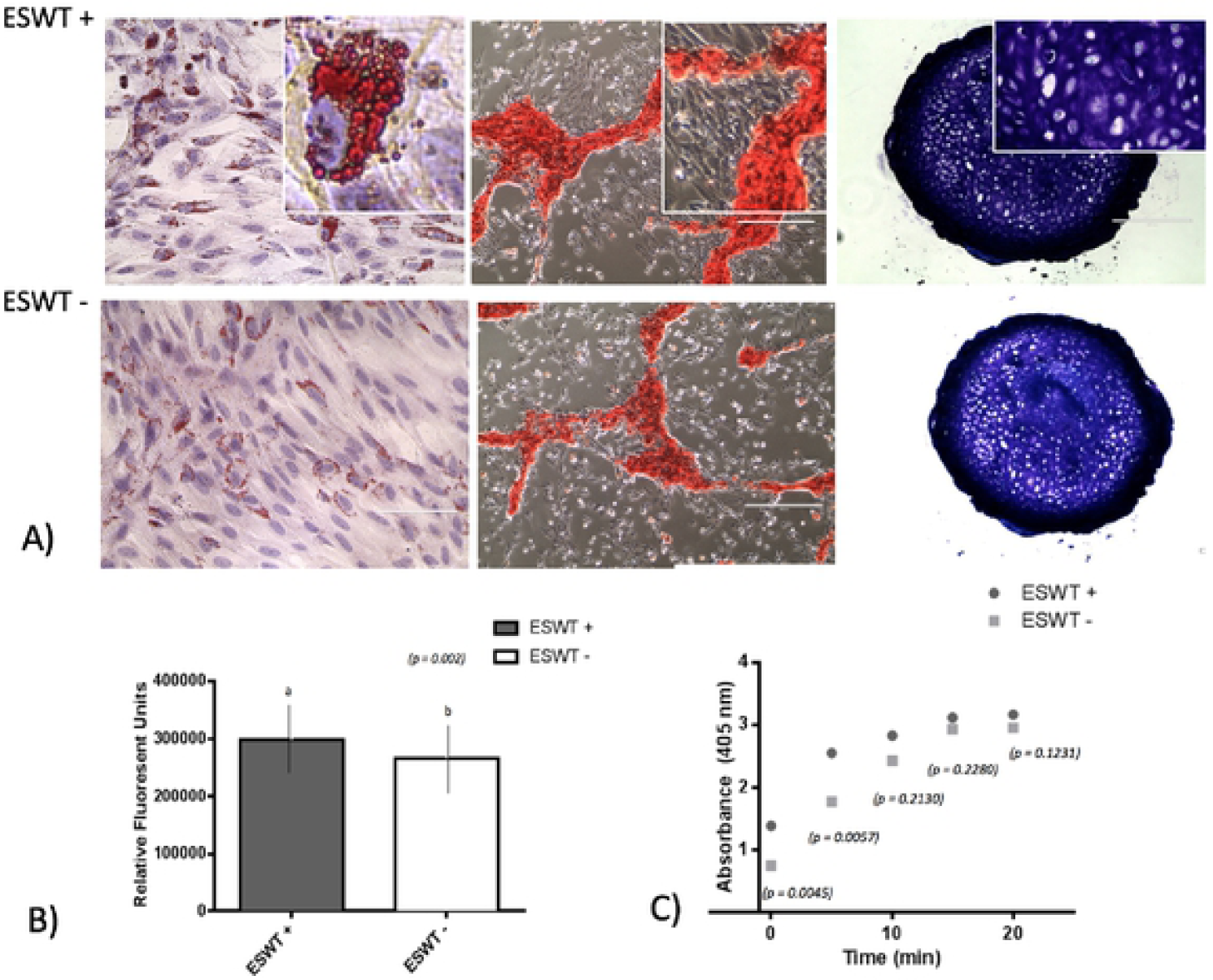
Trilineage differentiation. A) Positive trilineage potency for adipogencsis (Oil Red O), ostecogenesis (Alizarin Red S) and chondrogenesis (Toluidine blue) for treated (ESWT +) and untreated (ESWT -) equine CB-MSC. B) Quantification of intracellular lipid droplets by relative fluorescent units for treated (ESWT +) and untreated (ESWT -) equine CB-MSC. Results are shown as mean and CI 95%(*p* = *0.0002*). Values with different letters are statistically different (*p* = *0.002*). C) Alkaline phosphatase activity measured in relative fluorescent units in treated (ESWT +) and untreated (ESWT -) equine CB-MSC. Differences were found at time 0 (*p* = *0.0045*) and 5 minutes (*p* = *0.0057*).

### Mononuclear cell suppression assay

To investigate the immunomodulatory potency of equine CB-MSC treated with ESWT, we used a pooled mononuclear cell suppression assay (pMSA) method to compare MNC suppression of equine CB-MSC with and without ESWT. Treated and non-treated equine CB-MSC were co-culture with pooled MNCs for 96 hours and MNC proliferation was assessed with BrdU ELISA. Both treated and non-treated equine CB-MSC were able to suppress MNC relative to the positive control. The ESWT treated and non-treated MSCs suppressed MNCs similarily (Fig 7).

**Fig 7.**
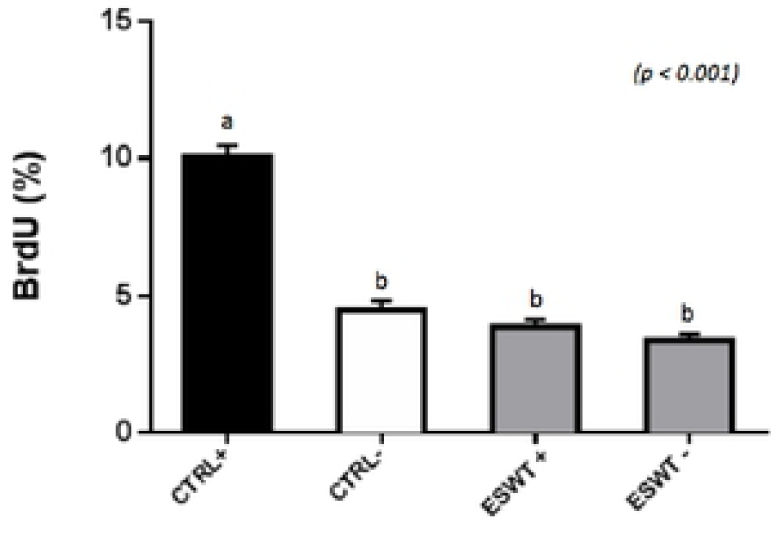
Mononuclear cell suppression assay. MNC suppression assay to determine MNC proliferation suppression capacity of equine CB-MSC. Both treated (ESWT +) and untreated (ESWT -) equine CB-MSC were able to suppress the proliferation of stimulated lymphocytes to the level of unstimulated lymphocytes (CTRL -). Values with different letters are statistically different.

## Discussion

In the present study, we demonstrated that equine CB-MSCs responds to ESWT stimulation. The proliferation of these cells was not adversely influenced by shockwave treatment but the metabolic activity of the treated cells was increased. Others have reported similar findings where the proliferation of human adipose-derived stem cells (hASCs) and rat adipose-derived stem cells (rASCs) were not different between the untreated control group and the ESWT group [8]. In contrast to those results, Suhr *et al.* observed that the application of ESWT to human bone marrow stromal cells (hBMSC) stimulated proliferation [7]. Weihs *et al*. showed an increase in both metabolic activity and in proliferation in a dose and time-dependent effect [10]. Varying treatment protocols, ESWT generators and application, species and stem cell sources could all account for the observed differences in proliferation and metabolic activity. MSCs isolated from different tissues possess a different ability to differentiate, different surface marker expression and proliferative capabilities [21,22]. Also, MSCs respond differently to the same stimulus, as observed when human adult and neonatal cells from the same individual were induced for neuronal transdifferentiation potential and showed different neuronal features [21]. Similarly, cells from the umbilical cord blood have shown increased proliferation and potency compared to bone marrow or adipose tissue [22]. As a result of this different response, a protocol utilizing the most appropriate cell source and cell type is essential to obtain the best response to treatment with ESWT.

In the current study, shock waves were applied via a water bath. The water bath was specifically designed to allow propagation of the shock waves after passing the cell culture, as it has been suggested that waves that are not propagated will get reflected at the culture medium and will disrupt the upcoming waves as well as the cell monolayer [18]. Conflicting information about the *in vitro* application of shock waves is published. Some authors recommend the application of the shock waves to flasks with an MSC cell monolayer in a water bath as we reported here; while others suggest applying shockwaves directly to tubes or flasks containing an MSC cell suspension through an interface (water-cushion, pigskin) [7,23]. The advantages of using the water bath in this study standardizes conditions, like temperature and distance, as well as being a model that simulates the *in vivo* conditions present during treatment with ESWT. One of the disadvantages of the method we used in our study, is that some detachment of the cells in the monolayer can occur, and furthermore, the cells located at the periphery might receive lower doses of ESWT [24].

The beneficial effects of ESWT on wound healing have been shown on different species (25). Previous studies reported an increased expression in shockwave treated tissues of various growth factors such as vascular endothelial growth factor (VEGF), transforming growth factor-beta 1 (TGF-B1) and insulin-like growth factor 1(IGF-1). The increase in growth factor levels is thought to stimulate healing by increasing neovascularization, fibroblastic activity and proliferation [25,26]. Other studies in horses have shown no difference in overall healing time but significant improved quality of healing by a decrease in granulation tissue [27]. Studies *in vitro* have shown an increase in migration of hASC, as shown by the scratch assay, where treated cells filled the scratch faster than the control [24]. We could not reproduce this finding in our study as we did not see a difference in migration across the scratch after shockwave treatment. This reported difference in biological behavior might be attributed to the distinct cell types or treatment protocols, as previously mentioned.

We have shown that equine CB-MSCs maintain their multilineage differentiation potential after treatment with ESWT. Treated cells do not only maintain but also increase their potency towards adipogenic and osteogenic lineage. There is growing evidence suggesting that ESWT has an effect on MSCs differentiation [7,9,10]. Until now, the only experiment assessing the response of equine MSCs to ESWT reported that shock waves stimulate a higher potential for differentiation towards adipogenic, osteogenic and chondrogenic lineages of equine ASCs [9]. In contrast, Suhr *et al*. found that the adipogenic differentiation was unaffected but the osteogenic and chondrogenic differentiation potentials were reduced in hBMSC [7]. Similar to our results, Leone *et al*. found an improved differentiation towards adipogenic and osteogenic lineages in tendon-derived stem/progenitor cells (hTSPCs) [28].

In the search of MSCs with an increased rate of survival after transplantation, Heneidi *et al*. isolated and characterized a new population of adipose stem cells (ASCs) named: multilineage differentiating stress enduring cells. The cells were isolated using severe stress conditions [29]. In our study after exposing the equine CB-MSCs to ESWT, an increase in adipogenic potency was observed in addition to an increase in metabolic activity. It is possible that ESWT can precondition the cells, allowing them a superior performance when transplanted. Furthermore, the combination of ESWT with ASCs has been shown beneficial, as observed in experimental studies of brain damage in rats where this combination of treatments reduces the effect of local and systemic inflammatory immune reactions, oxidative stress and apoptosis and also reduce damage to the mitochondria in the brain [30].

Several *in vitro* and *in vivo* studies have demonstrated the enhancement of osteogenic differentiations of different type of cells after ESWT by measuring BMP-2 expression, alkaline phosphatase (ALP) activity, and calcium deposits [31,32]. This is in agreement with our findings, as we observed an increase in ALP activity, which is a recognized biochemical marker of osteoblast activity [33]. Furthermore, it is believed that TGFB1 is responsible for the increased activity of osteoblasts [34].

Effects on differentiation and proliferation of different type of cells after ESWT can be explained by mechanotransduction, which is how cells respond to mechanical stimulation and convert these stimulations into biochemical signals that can alter growth factor expression and cellular adaptation [35]. The complete mechanism of action still remains unknown but there is evidence indicating that ESWT can stimulate the mitogen–activated protein kinase (MAPK) cascade [36,37]. As reported by Weihs et al., shock wave treatment initiates the release of ATP activating purinergic receptors that enhance proliferation by increasing ERK ½ signaling [10].

The contribution of the immunomodulatory ability of MSCs to tissue regeneration and healing has been long discussed. Using a mononuclear cell suppression assay (MSA), we observed that ESWT did not affect the mononuclear cell (MNC) suppressive potency *in vitro*. Our study is the first to examine the effect that equine CB-MSCs treated with ESWT has on MNC suppression. A previous study compared the surface markers and cytokine secretions of macrophages, and have demonstrated that the expression of classical pro-inflammatory M1 marker CD80 and COX2 decreased after ESWT, shifting the monocytes into the anti-inflammatory M2 phenotype. ESWT had also reduced the secretion of pro-inflammatory cytokines by M1 macrophages [39]. ESWT may influence MNC subpopulations when applied directly, but whether ESWT can affect MNCs indirectly through modifying MSCs still requires more investigation. It is possible that ESWT can influence MSC surface marker expression and cytokine levels, which would subsequently affect MNCs. However, this effect cannot be observed on the level of MSC induced MNC suppression. In addition, it is also possible that varying the energy level of the ESWT could change the outcome we have observed.

In previous experimental studies in horses *in vivo*, desmitis of the suspensory ligament or tendinitis of the superficial digital flexor tendon was induced with collagenase. Afterward, the injured and the control limb were treated with ESWT. The results showed improved healing of the treated limbs related to stimulation of fibroblasts, a greater number of mitochondria, with an increase in TGF b1, as well as improvement in their ultrasonographic appearance, reduced the size and increased neovascularization [40,41]. The positive responses to ESWT mentioned in the previous reports, together with our results, suggest the possibility to translate these findings into treatments for soft tissue injuries in horses, especially tendon injuries. However, the cells fate and biodistribution after intralesional administration remain to be described [42]. It has been shown that low percentages of MSCs are found after intra-lesional administration, and the MSCs can be found up to 90 days after injection but it has been difficult to prove that they integrate into the tissue [42,43].

Our study was a proof of concept in an *in vitro* model and the biggest limitation is that we did not investigate the effect of the combination of ESWT and equine CB-MSC *in vivo*. There are more limitations worth mentioning. First, the exact energy of ESWT that needs to be applied to equine CB-MSCs is unknown. Despite the fact that the study showed a significant increase in metabolic activity and increased adipogenic and osteogenic potential, the results are limited to the specific experimental settings selected here and should not be applied to clinical cases. Prospective clinical trials are needed to evaluate the effects of ESWT on different sources of MSCs *in vitro* and *in vivo*.

Also, the present study was purely descriptive and does not deal with the specific cellular response of equine CB–MSC to ESWT.

In conclusion, our findings indicate that ESWT can enhance metabolic activity and does not adversely affect the proliferation and immunomodulatory properties of equine CB-MSCs. For trilineage differentiation, we showed that ESWT allowed equine CB-MSCs to maintain their differentiation capacity, even more, treated cells showed an increased potency towards adipogenic and osteogenic lineages. The combined effect of these therapies remains to be seen, but the anecdotal clinical impression of improved healing after the combination of these two therapies can be partially explained through the increased metabolic activity and no effect on their immunomodulatory properties observed in this study.

Further *in vitro* and *in vivo* studies will be required to fully investigate the effects of ESWT on CB-MSCs and the mechanisms by which the combination of these treatments appear to improve healing of soft tissue injuries.

## Acknowledgements

We would like to acknowledge William Sears for his assistance with the statistical analysis.

